# Granule cells affect dendritic tree selection in purkinje cells during cerebellar development

**DOI:** 10.1101/2022.10.18.512799

**Authors:** Mizuki Kato, Erik De Schutter

## Abstract

This study investigates the interrelationship between primary dendrite selection of Purkinje cells and migration of their pre-synaptic partner granule cells during cerebellar development. During development of the cerebellar cortex, each Purkinje cell grows more than three dendritic trees, from which a primary tree is selected to develop further, whereas the others completely retract. Experimental studies suggest that this selection process is coordinated by physical and synaptic interactions with granule cells. However, technical limitations hinder a continuous experimental observation of multiple populations. To reveal the mechanism underlying this selection process, we constructed a computational model of dendritic developments and granule cell migrations, using a new computational framework, NeuroDevSim. Comparisons of the resulting morphologies from the model demonstrate the roles of the selection stage in regulating the growth of the selected primary trees. The study presents the first computational model that simultaneously simulates growing Purkinje cells and the dynamics of granule cell migrations, revealing the role of physical and synaptic interactions upon dendritic selection. The model provides new insights about the distinct planar morphology of Purkinje cell dendrites and about roles of the dendritic selection process during the cerebellar development. The model also supports the hypothesis that synaptic interactions by granule cells are likely to be involved in the selection procedure.

**Author Summary:** The mature structure of a Purkinje cell, the main neuron of the cerebellum, is composed of a large flat dendritic tree composed of a single or sometimes 2 primary dendrites. However, this neuron has multiple similar trees during development, and retracts most of them to select its primary tree. We aim to explore roles of this developmental process as either an important step to attain optimal morphology or merely a redundant step to be eliminated by evolution in the future. We especially focused on environmental interactions on Purkinje cells as criteria to select the dendritic tree, hypothesizing that developing an efficient network with other neurons is one of the main goals of neuronal morphology. We constructed and used computational models to investigate detailed physical interactions and communications between Purkinje cells and their environment. Comparisons of models suggest the role of the selection stage is to obtain favorable growth of primary trees.

## Introduction

The cerebellum is involved in coordinating motor functions as well as in cognition and emotion (1–4). Major neurons in this complex information processor migrate and maturate postnatally in mammalian brains. The cerebellum comprises only 10% volume of the whole brain but holds more than 70% of all neurons. Most of these neurons are cerebellar granule cells whose population seems too numerous to be counted accurately (estimated densities range from 500,000 (5) to 6,562,500 (6) cells per mm^3^ in mice, compared in (7). Despite such numerical predominance, the cerebellum is sometimes referred to as simpler than the cerebrum partially because output from the cerebellar cortex is composed of a single type of neuron, the Purkinje cell. Purkinje cells receive two types of excitatory inputs, from parallel fibers and from climbing fibers. Parallel fibers are the axons of granule cells, and climbing fibers come from the inferior olivary nuclei in the medulla. Purkinje cells project inhibitory outputs to cerebellar nuclei which also receive collateral input from afferents to the cerebellar cortex (climbing fibers and mossy fibers).

During the early postnatal development phase, each Purkinje cell grows multiple dendritic trees (Fig 1A1) and then selects a primary tree among them by retracting most of the newly grown dendrites (Fig 1 A2) (8–10). Meanwhile, a large population of granule cells migrates from the surface to the bottom of cerebellar cortex (Fig 1 B1 to B2). This cortex layer reconstruction by granule cells creates a highly crowded environment for the Purkinje cell to grow and retract dendrites.

**Fig 1.**
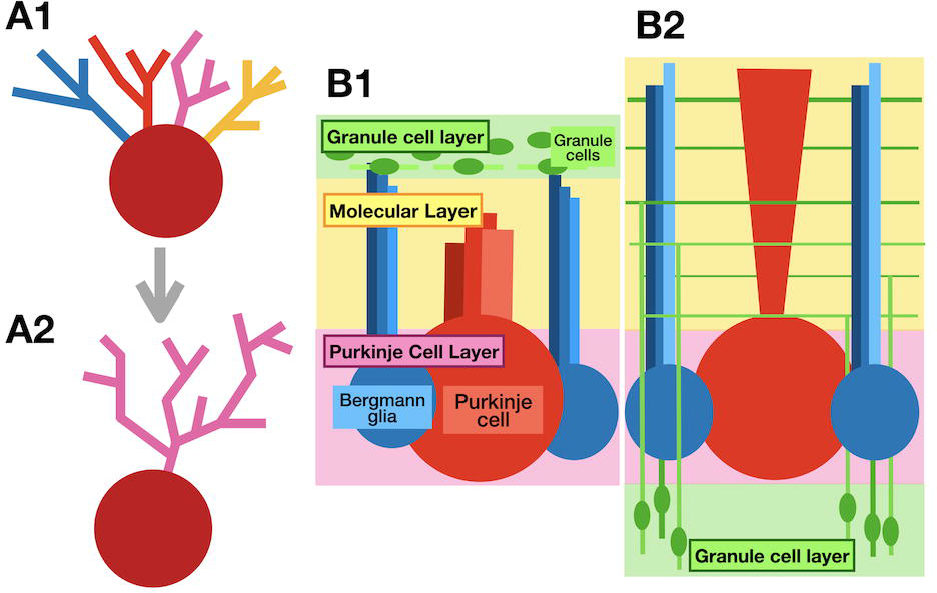
Schematics for dendritic selections of a Purkinje cell and cerebellar layer reconstruction by granule cells. (A1) A Purkinje cell before dendritic selection phase around postnatal day (P) 4 to 10. A red sphere represents its soma, and dendritic trees are shown in different colors. (A2) Same Purkinje cell after the selection phase around P8 to P10. In this schematic, the pink dendritic tree was chosen as its primary tree by retracting other candidate trees. (B1) The cerebellar cortex around P1, having granule cell precursors (green oval with light green lines as their leading processes or parallel fibers in the granule cell layer (shaded green) as an outermost surface. Its middle sheet is the molecular layer (shaded in yellow), and the bottom is the Purkinje cell layer. (B2) Structure of the cerebellar cortex after ^∼^P15. Granule cells (green ovals) migrated down to rearrange the granule cell layer (shaded in green) from the surface to the bottom. The molecular layer (shaded in yellow) comes to the surface and its volume expanded and is filled with parallel fibers and descending axons (green lines) extended by granule cells. The red structure represents a schematic Purkinje cell, and blue structures are Bergmann glia.

In isolated *in vitro* environments, Purkinje cells do not fully retract excessive dendritic trees, resulting in a state of multiple primary dendrites (11, 12). Granule cells are strong environmental factors to regulate the dendritic arborizations and retractions in terms of physical and synaptic interactions on Purkinje cells (13–16). However, involvement of the granule cells in the primary dendritic selection rule is still unclear.

The experimental approach to investigate the dendritic selection stage of Purkinje cells generates only a few samples of *in vivo* time-lapse images from early postnatal animals due to technical difficulties, therefore it is beneficial to extend the real samples with computationally constructed models.

Computational models simulating the interactions of granule cells and Purkinje cell populations during early postnatal days were built, and comparisons of models with varying parameters controlling the effect of intercellular interactions provided insights in how granule cells guide dendritic selection and arborizations in Purkinje cells.

## Results

### Granule cell migration model

In order to provide a 3D crowded environment for Purkinje cells to grow, a granule cell migration model was built first. The model simulates soma migrations and extensions of axons and parallel fibers of about 3,000 granule cells with additional 10,000 parallel fibers growing in from extended y-axis volumes (Fig 2A).

**Fig 2.**
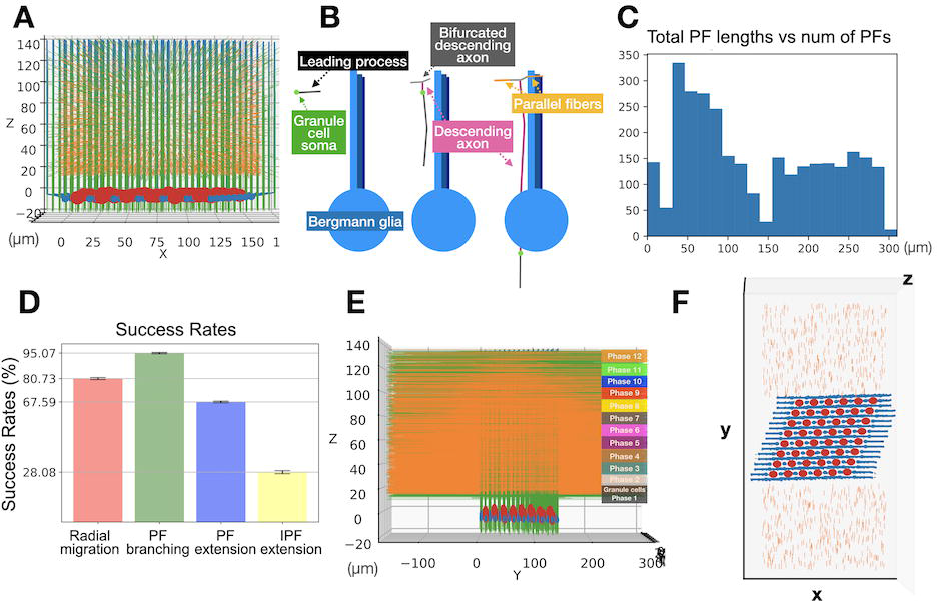
Granule cell migration model as an environment for Purkinje cell growth. (A) A representative simulation cube at the end of the simulation visualized from Y side. Red large spheres are Purkinje cell somas. Blue small spheres with processes growing upward are Bergmann glia. Numerous green thin threads are axons of granule cells in the main volume (where Purkinje cell somata and Bergmann glia are present), and orange fibers are ingrowing parallel fibers. (B) Schematic representations of how each granule cell migrates using a Bergmann glia process as a guide, see text for explanation. (C) A histogram showing final lengths of the parallel fibers in the representative case, 300 μm is the maximum possible length for the simulation volume. (D) Bar plots show mean success rates from 10 simulation samplings from the model with error bars showing standard deviations. A red bar is for success rate of soma radial migrations, green is for success rate of parallel fiber branching to form T-junctions, blue for parallel fiber extensions initiated in the main volume, and yellow for parallel fibers growing from the outside into the main volume and fully extending in the main volume. Coefficient of variation values for each is 0.031, 0.007, and 0.007, and 0.004 respectively. (E) Another side view (longitudinal side) of the granule cell model after all green granule cells finish migration and color stripes represent z-locations of granule cell initiation from 1 to 12 phases. (F) Top view of the granule cell model right after initiations of granule cells and incoming parallel fibers (at cycle = 8). Large red spheres are Purkinje cell somata, and smaller blue spheres are somata of Bergmann glia with their processes. Incoming parallel fibers are initiated at the extended volumes in y-axis. Thin orange cylinders in the extended areas are the ingrowing parallel fibers, and tiny black structures in the main area (where Purkinje cell somata and Bergmann glia line up) are horizontally migrating granule cells.

The sequences simulated during granule cell migration are shown schematically in Fig 2B. First, a granule cell soma (a small green dot) migrates horizontally to one of the closest Bergmann glia processes (light blue, more distant glia processes of same cell are dark blue) by extending a cylindrical leading process (gray). Next, the granule cell soma gets close enough to the target process, and starts radial migration by extending the leading process downwards along the process while extending an axon (pink) from the tail of soma which will further bifurcate into parallel fibers. The soma keeps migrating down and parallel fibers further extend along the y-axis in both directions. A line of axonal fronts lays down along the path of the radial migration of the soma.

In a representative model case, 80.59% of the granule cells succeeded in the radial migration, and 67.73% of parallel fibers extended through the complete volume of y-axis though most of them ended up with shorter lengths than experimentally observed (Fig 2C). Also, only 29.31% of incoming parallel fibers completed extensions through the whole volume. However, total parallel fibers in the central volume was 3487 towards front and 3430 to back. Distributions of success rates from 10 simulations are shown in Fig 2D.

## Results from 3 control Purkinje cell growth models

### Introduction

The goal of these 3 unique dendritic growth scenarios is to provide controls for scenarios that simulate specific retraction strategies. The control models specifically focus on investigating the influence of dendritic selection processes and physical presence of granule cells on dendritic tree morphology. We simulated growth up to P10. Two phases of retraction are implemented in all growth scenarios, which is inspired by unpublished statistical data of Purkinje cells from Prof. Yukari Takeo where the final dendritic selections *in vivo* usually turn out to be competition between two remaining candidate trees.

Simulations for scenario 0A and 0B used the saved initial conditions, and initiation of dendrite growth starts at cycle 65. The 0A_growOne scenario randomly chooses a single dendritic tree out of five candidates at cycle 65 + 7 and retracts all others (visible at the beginning in each plot in Fig 3 A1-A3). On the other hand, S0B_growAll scenario never initiates retractions and all the dendritic trees in each cell keep growing until they reached z = 100 μm (Fig 3 B1-B3).

**Fig 3.**
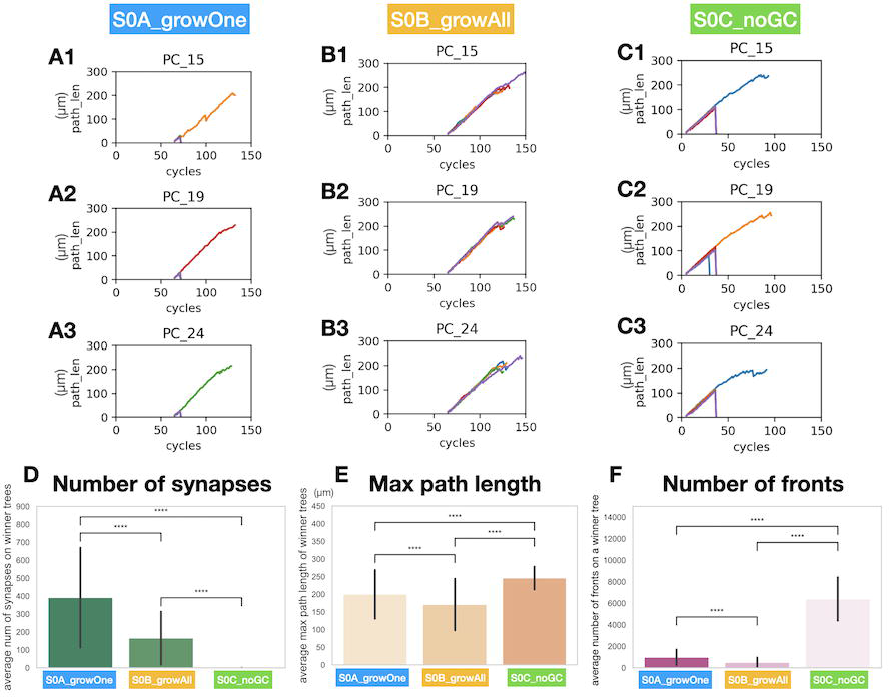
Characteristic of 3 control scenarios. (A1-3) The plots show change in max path length of each candidate tree at every cycle with a different line color for each dendrite. Three Purkinje cells were randomly selected from one of 10 simulations in scenario S0A growOne. (B1-3) Similar plots as A1-3 from one of 10 simulations in scenario S0B growAll. (C1-3) Similar plots as A1-3 and B1-3 from one of 7 simulations in scenario S0C noGC. (D) Plotting the number of synapses on each dendritic tree in the three scenarios with error bars representing standard deviations. (E) Plotting max length of each tree in the three scenarios with the error bars. (F) Plotting number of fronts in each tree in the three scenarios with the error bars. P values: **** indicates p < 0.00005

S0C_noGC scenario did not use the saved database as there are no granule cells at all, and dendritic growth is initiated at cycle 5. After cycle 30, the simulation checks if cells grew more than 250 dendrite fronts (^∼^5μm per a front) in total. If a cell reached that threshold, front numbers are computed for each dendritic tree, and a first retraction occurs of trees that have less than 60 fronts. At cycle 30 + 7, a second check proceeds, choosing the tree with the largest number of fonts for cells having at least 400 fronts in total and retracting all other trees. In the example cells in Fig 3 C1-C3, Purkinje cell 19 (PC_19, Fig C2) follow this two-phased retraction as dendritic trees are retracted in 2 different cycles. PC_15 (Fig C1) and PC_24 (Fig C3) retracted dendrites at a single cycle because all dendritic trees at the first screening phase were mature enough to have more than 60 fronts and retractions only initiated at the second phase.

### Number of synapses, max path length, and number of fronts

A comparison of the average number of synapses with parallel fibers in winner trees shows differences between scenarios S0A, S0B, and S0C (Fig 3D). Most clearly, dendritic trees in S0C did not have any synapses because there are no granule cells. Each dendritic tree in S0A_growOne got a much larger number of synapses than S0B_growAll, probably because growing all candidate trees led to more intense competition over limited numbers of synapse locations with parallel fibers. Moreover, in S0B, more structures in the simulation cube can lead to a decrease in the number of fully extended parallel fibers, which makes the competition even more severe. Although experimental data to compare number of synapses on dendritic trees at the age of interest is not available, it seems the model provided enough opportunity for dendrites to form synapses with parallel fibers. Assuming that there are enough parallel fiber fronts to form synapses, the larger max path length in S0A than S0B contributed to the increase in number of synapses (Fig 3E). Winner trees in S0C had an artificially large increased number of total fronts (Fig 3F) since, by default, the model uses smaller growth steps when no parallel fiber is detected, resulting in (more) smaller fronts.

### X-distance and y-distance

Characteristic morphology of the winner trees from these 3 scenarios was determined with further tree structure analysis (Fig 4). Especially, average max y-distance of the winner tree was greatly influenced by physical presence of granule cells (Fig 4 A1-C1, and E). This was due to a growth algorithm of dendritic tree which directs growth along a plane perpendicular to close by parallel fibers (see Methods). The algorithm included dendrite-dendrite repulsions, which also contributed to flatness of the tree. But these repulsions have to be combined with physical presence of granule cells to establish distinct planar morphology as in S0A and S0B.

**Fig 4.**
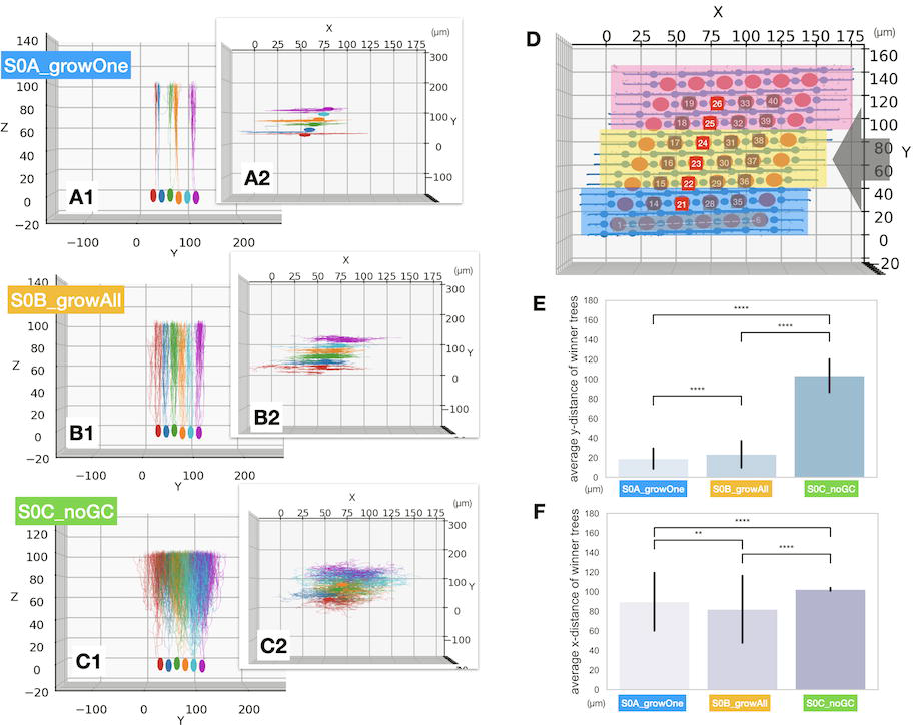
Arbor size of winner trees in 3 control scenarios. (A1) visualizations of Purkinje cells (pc_21 in red, pc_22 in blue, pc_23 in green, pc_24 in yellow, pc_25 in light blue, and pc_26 in magenta) from side view at cycle 120 of 220 for growth scenario S0A. Only selected Purkinje cells were visualized for convenience, while other structures (e.g. granule cells, Bergmann glia, and other Purkinje cells) are not shown. (A2) A top view of A1. (B1) A similar plot for scenario S0B. (B2) A top view of B1. (C1) A similar plot for scenario S0C. (C2) A top view of C1. (D) Top view of the original granule cell model to show the map of Purkinje cells visualized in A-C. The grey arrow indicates the viewpoints for A1, B1, and C1. (E) Plots for comparing average max y-distance of each tree from the three scenarios with error bars indicating standard deviations. F) Plots for average max x-distance for the three scenarios with error bars. P values: **** indicates p < 0.00005 and ** is p < 0.005

Compared to the results from max y-distance, no remarkable difference in max x-distance was found at a glance (Fig 4 A2-C2, and F). However, *P* values indicated significant differences between the data sets, and S0A and S0B has greater variation between samples compared with S0C. Physical hindrance from granule cells possibly contributed to extend the divergence in the tree morphology. Comparing the y-distance of S0A and S0B, S0A has slightly thinner trees than S0B. This was unexpected because competition over space availability in Y axis directions in S0B is more severe than S0A.

In 150-180 postnatal day mouse, y-distance of dendritic trees in control mice was 13.9 ±1.5 μm and x-distance was 82.9 ±2.5 μm (17). Y-distance from scenario S0A and S0B are slightly thicker than the observed morphology, though Purkinje cell dendritic trees attains single planar morphology only after postnatal day 22 in mice (18) which is later than up to postnatal day 10 simulated in the model. For growth in x-axis direction, the dendritic tree reaches its full x-width roughly by postnatal day 13 in mice (10). The x-distance of the 3 scenarios was marginally wider than that observed. Extended y-distance (28.7 ±4.8 μm) and shrinkage of x-distance (40.0 ±4.6 μm) was observed in reeler mice where the structure of cerebellar cortex is destroyed due to a failure of granule cell migration (17). S0C also got expanded y-distance compared to S0A and S02.

### Number of branch points and terminals

In addition to the tree size in x- and y-axis, dendritic morphology was further analyzed by comparing branch points and terminals of winner trees (Fig 5). In general, winner dendritic trees in S0C have much larger numbers of fronts, branch points and terminals.

**Fig 5.**
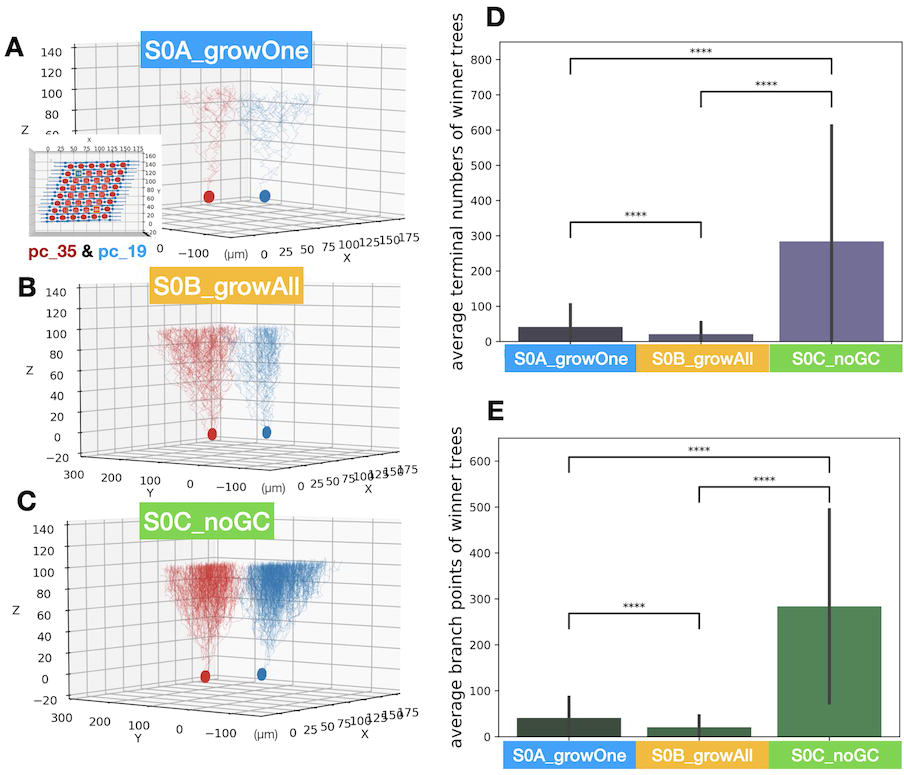
Morphology of winner trees in 3 control scenarios. (A) Two Purkinje cells (pc_35 in red and pc_19 in blue) at cycle 120 of 220 for scenario S0A are visualized. (B) The same plot as A for scenario S0B. (C) The same plot as A and B for scenario S0C. (D) Plots for comparing average numbers of terminals in each tree from the three scenarios with error bars indicating standard deviations. (E) Plots for average max x-distance for the three scenarios with error bars. P values: **** indicates p < 0.00005

The smaller number of branch points and terminals in S0B winner trees compared to S0A imply that multiple dendritic trees growing from the same cell reduces the elaboration of each tree. This assumption suggests that a role of the dendritic selection stage of Purkinje cell development may be to promote elaboration of a single tree to avoid intensive competition in space and synapse in the distal area, while leaving extra space and the resources in proximal area.

### Control growth scenarios: conclusions

The simulation results from these scenarios suggest that the extended granule cell model provided enough numbers of parallel fibers for Purkinje cells in the model. There was a small discrepancy in morphology of the area proximal to the soma compared to the real Purkinje cells, this was due to how the growth model was defined. However, these growth models suggest that excess trees growing from the same cell lead to a reduced dendritic tree morphology by competition for space and synapses, implying an important role for the retraction stage. Also, the comparisons show that the physical presence of granule cells contributes to the distinct planar structure of Purkinje cell dendritic trees.

## Results from 3 purkinje dendrite retraction scenarios

### Introduction

Two phases of retraction are implemented in all scenarios, timings and criteria of retractions depend on given parameter sets. The first scenario triggers the retractions by fixed timings so that all Purkinje cells retract at the same time. The second scenario triggers retractions by two types of maturation status: tree size or number of synapses, and the retractions happen at different times depending on maturation levels of each cell. The third scenario triggers retractions by value of integrated synaptic signals, but is otherwise similar to scenario 2. By comparing with control growth scenarios, the additional role of the dendritic retraction stage to enhance expansion of selected dendritic trees could be analyzed.

### Retraction parameter space evaluations

The number of synapses and path length of winner trees corresponding to different parameter sets in retraction scenarios 1 through 3 are shown in Fig 6 (A, B1, B2, and C). The size of the circles on plots represents max path length of the winner trees and had similar ranges for all scenarios.

**Fig 6.**
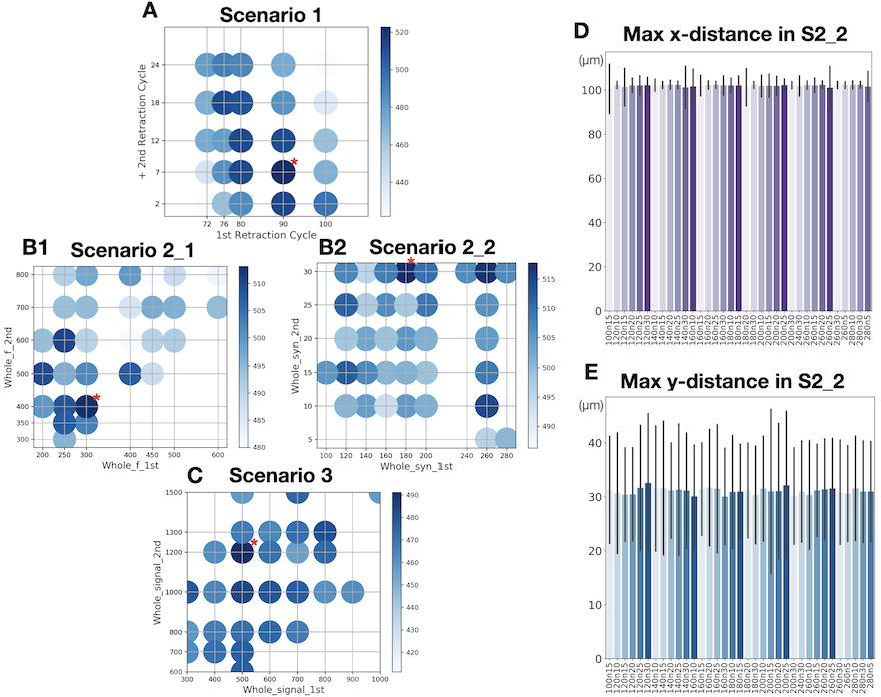
Dependence of number of synapses and arbor size of winner trees on parameter sets. (A) The organized scatter map shows number of synapses by depth of colors with each combination of parameters. As shown in color bar, circles have darker color for increasing number of synapses. X-axis represents the first retraction cycle and y-axis shows number of cycles after a parameter for the first retraction cycle. For example, actual 2nd retraction cycle is 72 + 2 = 74 if the sample’s 1st retraction cycle was 72, 2nd retraction cycle is 76 + 2 = 78 if its 1st retraction cycle was 76, and so on. The sizes of circles on the plot also represent average max path length of winner trees. The parameter set with the most numbers of synapses is marked with a red asterisk. (B_1) A similar plot as A for scenario 2_1, but x-axis is Whole_1st threshold for fronts numbers, y-axis is for Whole_2nd threshold, and 1st_f_th is fixed at 20. (B_2) A similar plot for scenario 2_2. X-axis shows Whole_2nd threshold for number of synapses, y-axis for 1st_s_th synapse threshold, and Whole_1st synapse threshold is fixed at 90. (C) Similar plot for scenario 3. X-axis for Whole_signal_1st threshold and y-axis for Whole_signal_2nd threshold. A and C used 10 simulation samples for each parameter set, but three parameter sets with the largest number of synapses used additional 10 simulations. B_1, and B_2 used 20 simulation samples for each, and those trees that got 0 synapses but stayed until the end of simulation were excluded. (D) Plotting average max x-distance of winner trees from one group of simulation sets in scenario 2_2 as a representative example for all scenarios. Vertical lines are error bars indicating standard deviations. (E) The same plot as D, but for average max y-distance.

There was a high variance in the number of synapses generated for different parameter combinations for all scenarios. Complete color maps of different parameter sets could be made for scenario 1 and 3. Scenario 2_1 by fronts and scenario 2_2 by synapses are controlled by 3 parameters, in Fig 6B1-B2 we show combinations of two parameters, with the third parameter fixed. While for scenarios 1 and 3 clear trends were visible in the color maps, this was less the case for scenarios 2_1 and 2_2 that resulted in noisier maps.

Highest average number of synapses on winners for scenarios (red star in Figure 6) were as follows:

scenario 1: 530.43

(WholeCheck_cycle_1 = 90, WholeCheck_cycle_2 = 90+7)

scenario 2_1: 513.17

(Whole_1st = 300, 1st_f_th = 20, Whole_2nd = 400)

scenario 2_2: 517.76

(Whole_syn_1st = 90, 1st_s_th = 30, Whole_syn_2nd = 180)

scenario 3: 491.18

(Whole_signal_1st = 500, Whole_signal_2nd = 1,200)

Data from these best parameter sets were further analyzed in the following Figures.

Scenario 3 had a lower average number of synapses than other retraction scenarios, because thresholds based on integrated synaptic input may sometimes select smaller trees. An important difference between the scenarios is that scenario 1 forces two phased retractions at fixed cycles, choosing first two winners and then the final winner based on relative numbers of synapses. In contrast, scenario 2_1, 2_2 and 3 start the first checking at cycle 90 and then the second one at cycle 97 (values determined in scenario 1) for number of fronts, number of synapses, and signals respectively, and then choose different retraction timings for every cell depending on their specific criteria.

### Max y-distance and max x-distance

Max x-distance and y-distance of winner trees were also compared, but all selected trees in all scenarios of different parameter sets resulted in similar morphologies in terms of the size (Fig 6D, E). As mentioned in the control scenario, x-distance of P120-180 mice is 82.9 ±2.5 μm and y-distance is 13.9 ±1.5 μm (17). Both x-distance and y-distance of winner trees in the retraction scenario is larger than experimentally observed, but they are still in acceptable range considering mouse Purkinje cell dendrites attain their final x-distance roughly by P13 in mice (10) and the y-distance around postnatal day 22 (18).

### Branch points and terminals

Number of fronts, branch points, and terminals from all retraction scenarios were strongly correlated with each other (Fig 7A1-3). However, results from scenario 3 had a slightly smaller tree which correlated to their smaller number of synapses (Fig 7B,C). Compared with control scenarios S0A_growOne and S0B_growAll, winner trees from retraction scenarios tend to have larger numbers of fronts and branch points, which suggests a role of the dendritic selection stage to elaborate winner dendritic trees.

**Fig 7.**
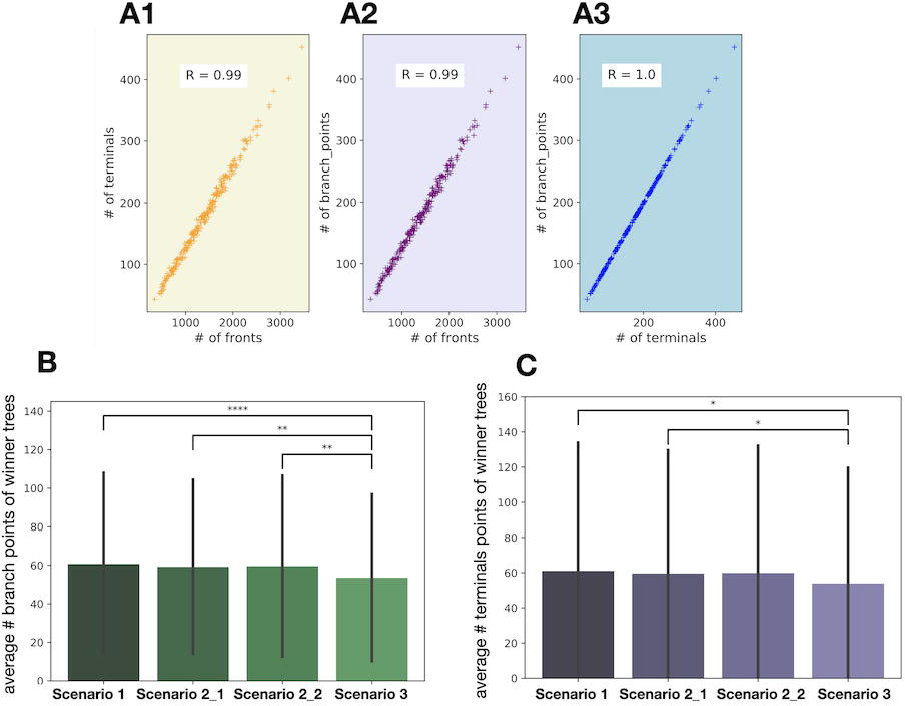
Morphology of winner trees in the retraction scenarios. (A1) Scatter plots show correlation between results from number of fronts and terminals for scenario 1, R is the correlation coefficient. (A2) A similar plot as A1 for correlations between number of fronts and branch points for scenario 1 results. (A3) A similar plot as A1 and A2 for terminals and branch points. Similar results were obtained for scenarios 2 and 3 (not shown). B) Plots for comparing average numbers of branch points on winner trees of scenario S1, S2_1, S2_2 and S3 with error bars indicating standard deviations. C) A similar plot as B for number of terminals. P values: * indicates p < 0.05 and ** is p < 0.005

### Retraction timings (path length vs. cycle)

When comparing individual data of retraction timings (Fig 8), Purkinje cells in all scenarios experience the second retraction at similar simulation cycles. As these represent optimal parameter sets to generate the largest number of synapses in each scenario, there may be an optimal timing to induce retractions to have more synapses although we do not have experimental data to compare with.

**Fig 8.**
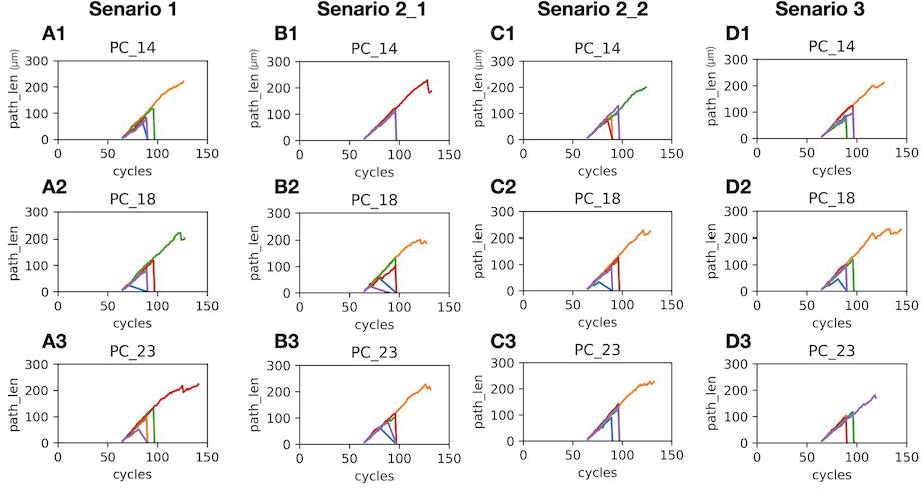
Change in path lengths with increasing simulation cycles in retraction scenarios. (A1-3) The plots show change in max path length of each candidate tree at every cycle with a different line color for each dendrite. Three Purkinje cells were randomly selected from one of 20 simulations in scenario 1. Since scenario 1 induce retractions at 2 fixed cycles, max path lengths of unselected trees dropped to 0 (retracted) at cycle 90 and 97. Two candidate trees in the sample (blue line on A2 and purple one on A3) could not fully grow as other trees and resulted in odd retraction lines. (B1-3) Similar plots as A1-3 from one of 20 simulations in scenario 2_1. All cells skipped the first retraction because all the dendritic trees had a larger number of fronts than the given threshold (20 fronts for this case) when a cell reached the whole cell front number threshold (300 fronts). Then, a winner was chosen when a cell reached 2nd whole cell front number threshold (400 fronts). (C1-3) Similar plots as A1-3 and A1-B from one of 20 simulations in scenario 2_2 whose 1st retraction thresholds for synapse numbers in each tree was 30 and in whole cell was 90, and whole cell threshold for 2nd retraction was 180 synapses. (D1-3) Similar plots as others but from one of 20 simulations in scenario 3 whose 1st retraction threshold for signal in whole cell was 500 and 2nd was 12,00.

### Distributions in number of synapses

In addition to the averaged data of multiple simulation samples, randomly selected individual synapse data and morphology of winner trees in single samples of each scenario are presented in Fig 9. Distributions in number of synapses from all samples of the retraction scenarios are shown in Fig 10. There were obvious differences between the scenarios in the range of number of synapses in the winner trees. Scenario 2_1 clearly has a much lower variability (Fig 9B1), possibly due to selecting winners by tree size which may balance out structural differences. For other scenarios the distribution is different with scenario 1 (Fig 9A1) and 2_2 (Fig 9C1) showing mean trends that are less obvious for scenario 3 (Fig 9D1); scenario 3 also has the largest variability overall. Randomly selected Purkinje cells from the same simulation as Fig 9 A1, B1, C1, and D1 are shown in A2-3, B2-3, C2-3, and D2-3. It is difficult to detect the morphological differences among each scenario from the figures, but varied morphologies of Purkinje cell dendrites from the same simulations can be seen. Movies for the dendritic growths and retractions of selected cells in each scenario can be found in supplementary materials. Number of synapses in all winner dendritic trees at cycle 140 for each scenario is also shown in Fig 10. Statistical differences of each data set by *p* values are shown at the bottom of Fig 10. Purkinje cells in scenario 3, maybe the physiologically most realistic model among all the scenarios, had the characteristic of sometimes choosing trees with relatively smaller numbers of synapses as their primary dendrites, resulting in a larger overall variability compared to other scenarios.

**Fig 9.**
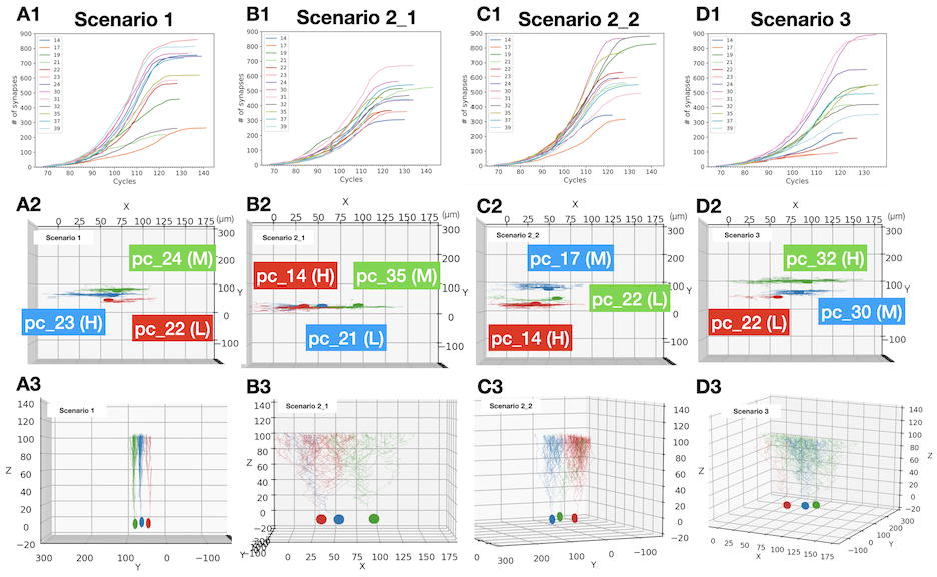
Changes in synapse numbers and morphology of individual winner trees from single samples in the retraction scenarios. (A1, B1, C1, and D1) Plots to show change in number of synapses at every simulation cycle in one of the simulations (seed 11) from the parameter sets that generated largest numbers of synapses in each scenario. (A2, B2, C2, and D2) Top view of selectively visualized Purkinje cells at cycle 220/220. Purkinje cells with highest (H), lowest (L), and somewhere in middle (M) number of synapses from A1-D1 plots were selected for each. (A3, B3, C3, and D3) Plots to show morphology of winner trees in A2-D2 from different directions. Plotted Purkinje cell samples are selected randomly, and numbers in legend corresponds to numbers in Purkinje cell location map on Fig 4D. Scenario 1 data is from a seed 1 simulation of a parameter set with WholeCheck_cycle_1 = 90 and WholeCheck_cycle_2 = 90+7. Scenario 2_1 data is from a seed1 of a parameter set Whole_1st = 400, 1st_f_th = 20, and Whole_2nd = 500. Scenario 2_2 is from seed 1 of Whole_syn_1st = 90, 1st_s_th = 30, and Whole_syn_2nd = 160. Scenario 3 is from seed 1 of Whole_signal_1st = 10,000 and Whole_signal_2nd = 13,000.

**Fig 10.**
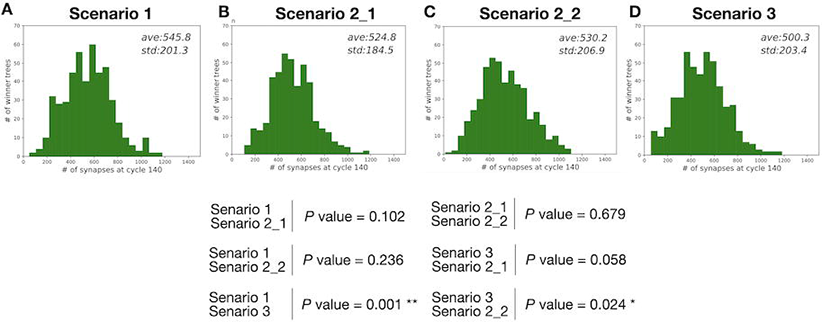
Variability of synapse numbers in all winner trees for the retraction scenarios. Histograms for numbers of fronts on winner trees from all 20 simulations in each scenario. Average and standard deviation of each data set is shown in each plot. P values for comparing each data set are listed at the bottom where * indicates p < 0.05 and ** is p < 0.005.

### Retraction scenarios: conclusions

When compared with tree morphologies from control scenarios, winner trees from retraction scenarios tend to have a larger number of fronts and branch points that suggest a role of the selection process to promote the growth of primary dendritic trees.

In terms of having opportunity to form more synapses, common optimal timings of retractions were observed across all retraction scenarios. The comparisons of scenarios by two different definitions of maturations (scenario 2_1 by size vs. scenario 2_2 by synapses) showed that choosing winners by sizes of the trees might not be the best choice compared to directly choosing winners by the largest number of synapses

Also, when the selection was controlled by integrated synaptic signals (scenario 3), Purkinje cells started to favor sometimes less elaborated trees with smaller numbers of synapses as their primary dendrites. Early born parallel fibers extending in the bottom of the molecular layer mature earlier, increasing their average firing rates. These stronger signals in the bottom of the layer may neutralize the advantage of having larger dendritic trees with more synaptic partners located in upper regions of the simulation volume.

## Discussion

### Summary of the main results

A Purkinje cell growth model with continuous growths of all dendritic tree candidates on the same cell suggested that dendrite-dendrite repulsion may not be the main factor to keep its unique flat structure. Also, the Purkinje cell models suggested possible roles of the dendritic selection stage in facilitating sophistication of trees and in balancing a consistent elaboration of the vertical area of trees. Meanwhile, keeping multiple dendritic trees until later developmental stage may result in smaller winner trees.

The model then explored detailed mechanisms of dendritic selection using 3 scenarios. While for retraction scenarios 1 and 3 clear trends were visible in the parameter maps for number of synapses in winner trees, this was less the case for scenarios 2_1 and 2_2 that were noisier. This may be due to the use of a fixed threshold for screening candidate trees at the first retraction stage. Using a relative threshold scheme, at least for the first retraction phase, might be more optimal for finding most efficient combinations of parameter sets to earn largest number of synapses. Purkinje cells may therefore use relative thresholds to select the winner among their candidate trees rather than using shared, fixed ones. Using the size of arbor as criteria of the winner selection did not necessary result in obtaining a larger number of synapses with parallel fibers. Rather, number of synapses itself may be the more important factor to win the selection. This trend matches the biological rule that the selection is not size dependent (personal communication, Prof. Yukari Takeo and Prof. Michisuke Yuzaki). Conversely, synaptic input reflecting network activity was less efficient as selection criteria because it sometimes favors dendritic trees with smaller numbers of synapses.

### Limitations of the models

The largest difference between the models and biological systems is that cerebellar expansion during development was not implemented because volume expansion cannot be simulated in NeuroDevSim. The cerebellum undergoes a massive expansion, especially along the anterior-posterior plane (x-axis in the simulation cube) (19) during 2 weeks after birth in mice (20). While the anterior-posterior plane in embryonic days 17.5 mouse has half the length of its molecular layer axis (z-axis in the cube), the anterior-posterior axis in postnatal mice expands 7.8 times longer than the molecular layer which also expands significantly (19). Therefore, this lack of volume expansion strongly affects crowding of cells, making it more difficult to manage migrations and growths of large cell populations.

There is also a difference in the process of extending parallel fibers during granule cell migration between the model and real neurons. In murine cerebellum, parallel fibers already extend during horizontal migration. However, the model extended parallel fiber axons after horizontal migration. The purpose of horizontal migration in biological system is unclear, but it might be related to granule cell allocation upon expansion of the cortex (21), which is not implemented in the model. In the real cortex, it may be advantageous to already extend axons to preserve space for axonal expansion since the space just above horizontally migrating granule cells is filled with layers of proliferating granule cells, and it may be quite difficult to squeeze in to form new T-junctions. Since the model does not implement proliferations of granule cells, granule cells have free space to grow T-junctions above them, and consequently the model had quite a high success rate of T-junction formations.

For Purkinje cell models, apoptosis and reorganizations of soma positions in Purkinje cell layer were not simulated. Also, spiny dendrites and stem dendrites are not distinguished in the model. Climbing fibers and mossy fibers, two afferents to the cortex forming synapses with developing Purkinje cells (22, 23) are not included in the model. Climbing fibers are involved in organizing the final shapes of Purkinje cell dendrites (24). Also, only a reduced branching pattern in dendritic arborizations of Purkinje cells is observed in rats when climbing fibers were removed by thermocoagulation *in vivo* (25). Similarly, transient contacts of mossy fibers with developing Purkinje cells are likely involves in assembly of zonal circuit maps of the cerebellum rather than dendritic selections (26, 27). However, there is no strong evidence that they are not involved in early dendritic development, and inclusions of these axons may be considered in future models. Finally, development was simulated only until P10, excluding later growth of the winning dendritic trees from analysis.

### Potential improvements and future directions of the models

In the model, all granule cells are homogeneous in that there is no zonal patterning by molecular phenotypes or birth orders. Such inhomogeneity in granule cells can be represented by, for example, 2 different types of molecular diffusions in the model to facilitate the zoning. Also, we can set different types of granule cell objects and assign different preferences in making synapses with distal or proximal regions of Purkinje cells ‘dendritic trees as observed (28, 29). It is technically also possible to include granule cell proliferation, but in the absence of tissue expansion in the model this did not seem useful.

For the dendritic structure of Purkinje cells, an additional growth algorithm to facilitate more elaboration of the proximal region of dendritic trees should be introduced to match the observed morphology. Also, the model needs to use a thicker outer wall of Purkinje cells to prevent edge effects since x-width of dendritic trees grow beyond the internal area of the Purkinje cell cluster.

Influence of dendritic trees growing from the same cell on morphology of the winning dendritic tree was found in the model. There are interesting cases in retraction scenario 2 simulations, where some cells ended up with 0 dendritic trees due to too high thresholds, and having such devoid cells boosted the average number of synapses of winner trees in surrounding cells. These results were treated as outliers in the analysis, but it would be interesting to check specific influence of surrounding dendritic trees as environmental parameters separated from influence of dendritic trees from the same cell. Simulating later stages of development is also a compelling challenge.

The comparison of different retraction scenarios suggests that the 1st retraction threshold is more likely to be specific for each cell. It is worth to confirm that the threshold for 2nd retractions works similarly.

Moreover, the model assumed 2-phased dendritic retractions recognized by Prof. Yukari Takeo (personal communication), and it also interesting to compare models with uni-phase or multiphase retraction schemes to explore advantages of having the 2-phases retraction system.

## Contributions of the models

Experimentally inducing no granule cell environment in the cerebellar cortex will result in too many effects on its development, like disturbing tissue expansions, molecular interactions, or development of neurons other than Purkinje cells. Our models enabled us to observe the physical interactions of granule cell migrations and dendritic development of Purkinje cells without such constraints. This provided a clue for understanding the well-known yet unexplained flat dendritic structure as being shaped by physical constraints of granule cells.

Also, one of our dendritic retraction models reflected the experimentally observed size independence for the primary tree selections. This indicates an involvement of the synaptic signal transductions from granule cells in the selection process of primary trees in Purkinje cells.

Our computational work can be combined with experimental approaches to further explore relationship between number of synapses and size of the dendritic trees. Experimental explorations on young Purkinje cells are still lacking, and it may be technically possible to mark synapses on the dendrites and count them using *in vitro* microscopy. Then, we can check for correlations between the number of synapses and size of the dendritic trees. This can also be combined with *in vivo* observations to obtain the data at specific timing of the developmental phase such as right after the dendritic selections. In addition to the number of synapses, we can collect data about synaptic activities in the same manner by checking synaptic properties and compare them with the simulation data from the model. However, such experimental procedure is only plausible if synaptic plasticity is present in the young dendritic trees and if there is a reliable anatomical marker. Moreover, if tissue culture systems of early postnatal cerebellum would be possible, we can test for signal activities *in vitro* like selectively giving signals on one of the candidate trees and observe its effects on the selection.

Lastly, the model is the first attempt to reconstruct development of the main neurons in the cerebellar cortex with detailed interactions between different types of neurons. The real-life cerebellum has evolved to be very complicated, therefore revealing this complexity by experimental procedures is obviously important. Modeling helps to re-organize and optimize the numerous information gained from live samples and facilitates to understand the bigger picture of how things work.

## Materials and Methods

The NeuroDevSim software (30) was used to construct and simulate the models (https://github.com/CNS-OIST/NeuroDevSim). NeuroDevSim is a computational framework specialized in simulating phenomenological and 3D structural interaction with the environment for developing neuronal morphology. Simulations were executed on the high performance computing cluster Deigo at OIST, using AMD Epyc 7702 CPUs with 128 cores and 512GB memory. 64 cores are used for each simulation, and it takes about 3 hours to finish a single simulation. Python scripts for each model can be found in the Model Database (http://modeldb.yale.edu/267591, access code:1601003).

The model embodies a Purkinje cell layer and a molecular layer. Its Purkinje cell layer has somas of Purkinje cells and Bergmann glia. The molecular layer has simplified Bergmann processes for guiding granule cells which gradually fill up the layer with their descending axons and parallel fibers (Figure 1B). Simulation of the granule cell migrations are performed in a (x_1_, y_1_, z_1_), (x_2_, y_2_, z_2_) = (−20, - 160, −20), (180, 300, 140) μm^3^ volume, representing a part of a murine cerebellar cortex (lobule VI) during postnatal days 0 to 10. The x-axis of the cube represents sagittal plane, y-axis represents transversal plane (long axis of folium), and z-axis is the depth of the cerebellar cortex (Fig 2E,F). The volume of the cube is fixed, therefore cerebellar tissue expansion is not implemented in the model.

### Granule cell migration model

About 3,000 granule cells are gradually initiated in 12 consecutive phases in the central volume (y-axis −20 to 160, Fig 2E) along with incoming parallel fibers of 10,000 granule cells located outside this central volume (orange structures in Fig 2F). Each phase initiates around 250 granule cells in the main volume and about 850 in-growing parallel fibers at the extended y-axis volumes. Upon initiation, granule cells horizontally migrate to a nearby Bergmann glia process (Fig 2B) and start radial migrations after they get close enough to the process (Fig 2B). During the radial migration, each granule cell soma extends a descending axon which further bifurcates as parallel fibers (Fig 2B). The cells keep migrating until they reach the internal granule layer (z = −15μm), while parallel fibers extend up to the ends of y-axis (Fig 2B). Granule cells start with a low firing rate that increases when they arrive in the internal granule layer, each cell’s rate is chosen randomly. The incoming parallel fiber structures simulate only parallel fibers themselves, just a soma with fibers extending along the positive or negative y-axis.

Granule cells in the model often experience collisions with surrounding structures during their migrations and extensions of the axonal fibers. Built-in methods in NeuroDevSim are used to manage collisions; providing a bypass (*solve_collision*()) for migration, and finding an alternative space around a colliding structure (*alternate_location*()) for the fiber extensions. Also, the granule cell soma radius is modeled smaller than the actual value since physical shoving between cells cannot be simulated in the model, the smaller diameter representing the ‘squeezed’ soma structure of actual cells.

### Purkinje cell model

The Purkinje cell growth in the model corresponds to Purkinje cells in a mouse during postnatal days (P) 4 to 10, which starts by growing new dendrites on the upper sphere of each soma and ends by developing young dendritic trees followed by dendritic selections (Figure 1A).

The initiation of dendritic growths waits till early granule cell migrations are completed, and starts at initiation of phase 6 granule cell migrations. 10-20 simulations of the early granule cell migration phases were saved and used as the common initial conditions for the different Purkinje cell growth and retraction scenarios.

### Growth algorithm

The basic growth algorithm simulates growth of a tip of dendrite towards a nearby synapse free parallel fiber segment. Once it gets close to the target, it makes a synapse and continues to grow perpendicular to the parallel fiber as observed in biological systems (31). If no free parallel fiber segment is nearby, the dendrites continue growth along the direction of their current heading with a small upward force assuming phenomenological involvement of interneurons in the molecular layers (32). When dendrites experience collisions, the code use the built-in *solve_collision*() method to find a detour to reach a destined position. Repulsion force by the closest dendritic tree of other cells or of the same cell also affect the growth direction, with strength of repulsion depending on a distance along the y-axis to the closest dendrite segment. During the elongation process, random branching events occur with a small probability. A dendrite tip also initiates a branch towards its target parallel fiber if the direction to it strongly deviates from its current heading direction.

### Retraction Scenarios

To select the surviving dendrite, three different retraction algorithms are implemented in the model; trigger dendritic retractions at fixed cycles, by dendrite maturation status, or by relative network activity levels. The fixed timing scenario uses two fixed simulation cycles to trigger retractions. At first cycle, 3 out of 5 trees are retracted and later the 2 remaining compete to become winner(s). Winners are determined by relative numbers of synapses per dendritic tree in each cell. The second scenario initiates retractions by comparing two kinds of maturation levels; size of dendritic tree or number of synapses. Maturation by tree size focuses on an internal parameter of growth, and granule cells merely acts as physical obstacles. Conversely, maturation by synapses focuses on intercellular communication, and defines a tree maturation by number of synapses with parallel fibers. The third scenario, retractions for low network activity, focuses on synaptic interactions with dendritic trees and granule cells. Synaptic activity depending on the firing activity of granule cells is integrated into a *signal* and total *signal* activity in the dendritic tree is used as selection criterion.

### Control scenarios

In the first control scenario, 0A, Purkinje cells skip the dendritic selection stage by randomly selecting a single primary tree for each Purkinje cell 7 cycles after initiating growth of 5 candidate trees. The second control scenario 0B never triggers retractions of the candidate trees resulting in a physically more competitive environment for space availability. Physical hinderance by granule cell structures affect dendritic tree morphology in both scenario 0A and 0B. On the other hand, Purkinje cells in scenario 0C grows without granule cell migrations. The retraction mechanism for this scenario used the best parameter set from the retraction scenario 2 by tree size so that they can initiate retractions without granule cells.

### Data analysis

24 Purkinje cells enclosed by outer Purkinje cells were sampled for analysis. For all scenarios, morphology of resulted dendritic trees was analyzed and compared according to number of winner trees, and size, number of synapses, max y- and x-distance, and number of branch and terminal points of winner trees.

## Acknowledgements

This project is supported by the PhD program of Okinawa Institute of Science and Technology Graduate University (OIST). We thank the Scientific Computing and Data Analysis section of Research Support Division at OIST for providing the high-performance computing service for simulating our models and Prof. Michisuke Yuzaki at Keio University and Prof. Yukari Takeo at Stanford University for sharing their invaluable data.

## Supporting information

All movies show only a few Purkinje cells: cells 21 in red, 24 in blue, 29 in green, and 33 in yellow (see Purkinje cell location map on Fig 4D). Granule cells, parallel fibers and Bergmann glia are not shown. For each Purkinje cell, the dendritic trees in color are final primary trees, while ones that will be retracted are drawn in black. The movies start from simulation cycle 61 and ends at cycle 140.

**SI 1. Dendritic growths and retractions movie for control scenario 0A (grow one tree)**. Seed 2 of scenario 0A simulation was used to generate the movie.

**SL 2. Dendritic growths and retractions movie for control scenario 0B (grow all trees).** Seed 1 of scenario 0B simulation was used to generate the movie.

**SL 3. Dendritic growths and retractions movie for control scenario 0C (no granule cells).** Seed 1 of scenario 0C simulation was used to generate the movie, this movie starts from simulation cycle 0 and ends at cycle 140.

**SI 4. Dendritic growths and retractions movie for retraction scenario 1 (fixed timings).** Seed 1 of retraction scenario 1 simulation was used to generate the movie.

**SI 5. Dendritic growths and retractions movie for retraction scenario 2_1 (maturation by size).** Seed 1 of retraction scenario 2_1 simulation was used to generate the movie.

**SI 6. Dendritic growths and retractions movie for retraction scenario 2_2 (maturation by synapse).** Seed 11 of retraction scenario 2_2 simulation was used to generate the movie.

**SI 7. Dendritic growths and retractions movie for retraction scenario 3 (maturation by network activity).** Seed 11 of retraction scenario 3 simulation was used to generate the movie.

